# PAC-mediated AKI protection is critically mediated but does not exclusively depend on cell-derived microvesicles

**DOI:** 10.1101/2020.04.23.057745

**Authors:** H. Dihazi, K. Schwarze, S. Patschan, G.A. Müller, O. Ritter, D. Patschan

**Author notes:** **Contact information:** D. Patschan, MD PhD Medizinische Hochschule Brandenburg Klinikum Brandenburg Hochstraße 29 14770 Brandenburg.

## Abstract

Acute Kidney Injury (AKI) significantly worsens the prognosis of hospitalized patients. In recent years, cell-based strategies have been established as reliable option for improving AKI outcomes in experimental AKI. Own studies focused on so-called Proangiogenic Cells (PACs). Mechanisms that contribute to PAC-mediated AKI protection include production / secretion of extracellular vesicles (MV - microvesicles). In addition, the cells most likely act by paracrinic processes (secretome). The current study evaluated whether AKI may be preventable by the administration of either PAC-derived MV and / or the secretome alone. AKI was induced in male C57/Bl6N mice (8-12 weeks) by bilateral renal ischemia (IRI - 40 minutes). Syngeneic murine PACs were stimulated with either melatonine, Angiopoietin-1 or -2, or with Bone Morphogenetic Protein-5 (BMP-5) for one hour, respectively. PAC-derived MV and the vesicle-depleted supernatant were subsequently collected and i.v. injected post-ischemia. Mice were analyzed 48 hours later. IRI induced significant kidney excretory dysfunction as reflected by higher serum cystatin C levels. The only measure that improved AKI was the injection of MV, collected from native PACs. The following conditions worsened post-ischemic renal function even further: MV+Ang-1, MV+BMP-5, MV+melatonin, and MV+secretome+Ang-1. Together, our data show that PAC-mediated AKI protection substantially depends on the availability of cell-derived MV. However, since previous data showed improved AKI-protection by PACs after cell preconditioning with certain mediators (Ang-1 and -2, melatonine, BMP-5), other than exclusively vesicle-dependent mechanisms must be involved in PAC-mediated AKI protection.

## Introduction

In-hospital incidences of AKI (Acute kidney injury) have been increased over the last 10-15 years. In central Europe, it is being estimated that nearly 15% of all subjects treated in hospitals acquire AKI during the course of the disease [1]. The prognosis however has not significantly been improved since the early 1990s, particularly high mortality rates have been reported in AKI patients with malignancies, sepsis and dialysis [2]. Thus, clinical and experimental researchers all over the planet strive to identify new strategies for AKI diagnosis and management. Cell-derived therapies have been utilized in experimental AKI, the results are promising. Regarding Mesenchymal Stem Cells (MSCs), even some clinical trials have been initiated [3]. Another emerging cell population are so-called induced Pluripotent Stem Cells (iPSCs) [4]. Studies on iPSCs in AKI are rare at the moment and exclusively restricted to animal models [5,6]. Several own investigations focused on Proangiogenic Cells (PACs), which for many years were defined as early Endothelial Progenitor Cells [7]. Systemic PAC administration significantly attenuated murine AKI, in both terms, excretory function and renal morphology [8–10]. There are however certain problems associated with cell-based therapies in AKI in general. Firstly, cell isolation and expansion potentially require several days. Secondly, the exact time point of cell injection remains difficult to predict. Finally, cells, isolated from a certain donor must be accepted by an individual suffering from AKI. Therefore, cells, whatever their exact biological characteristics may be will most likely not be utilized for AKI treatment in a direct manner in the near future. The problems associated with cell-based therapies have been discussed lately [11].

Meanwhile, the mechanisms by which PACs act within the post-ischemic microenvironment have been elucidated more in detail. In 2012, Cantaluppi and colleagues [12] identified cell-derived microvesicles (MV) to be of critical importance for AKI protection. The administration of MV alone substantially protected animals from AKI. Several years earlier, Rehman et al. [13] identified a heterogenous group of humoral factors produced and secreted by PACs (EPCs). It was subsequently argued that a particular group of proteins, described by the term *secretome*, potentially mediates vasomodulatory effects of the cells. Today, the exact roles of MV and secretome in tissue protection are still unknown. We therefore aimed to analyze the effectiveness of both components in murine AKI. The overall goal was to determine which cellular excretory product may possibly be utilized for therapeutic purposes in the future.

## Results

Before the results will be presented, it needs to be stated that significant adverse events were not observed in any of the experimental groups. Therefore, no animals required euthanization before all analyzet were collected.

### PAC-induced AKI protection is mediated by MV, derived from native cells

Control mice displayed mean serum cystatin C levels of 342 ±20 ng/ml. Kidney function significantly deteriorated post-ischemia (696 ±26 ng/ml, p<0.0001). Three distinct treatment strategies were defined: administration of either the secretome (SECR) or microvesicles (MV) alone or of SECR and MV combined. Both components were either isolated from native cells or from stimulated PACs. Since previous own studies showed substantial PAC agonistic effects of Angiopoietin-1 / −2 [15,16], Bone Morphogenetic Protein-5 [17], and Melatonin [8], we decided to utilize each of these four mediators in an individual series of experiments. Therefore, we performed a total number of 15 interventional groups. Whereas native SECR or SECR / MV in combination did not improve kidney function post-ischemia (671 ±26 and 630 ±25 vs. 696 ±26 ng/ml; p=0.5 and p=0.08), MV derived from unstimulated cells significantly lowered serum cystatin C levels (593 ±17 vs. 696 ±26 ng/ml; p=0.004). Angiopoietin-1 (Ang-1): PAC preconditioning with Ang-1 for 1 hour did not improve therapeutic effects of SECR or MV. On the contrary, if MV from stimulated cells were applied, kidney function declined even further (IRI+MV+Ang-1 and IRI+SECR+MV+Ang-1 vs. IRI 922 ±34 and 875 ±28 ng/ml vs. 696 ±26 ng/ml; p<0.0001 and p=0.0002). Angiopoietin-2 (Ang-2): Ang-2 stimulation did not result in any improvement or further deterioration of post-ischemic kidney function. The numerical results shall not be given in the text (see Figure 1). Bone Morphogenetic Protein-5 (BMP-5): PAC stimulation with BMP-5 aggravated AKI if MV were applied alone (IRI+MV-BMP-5 vs. IRI 799 ±32 vs. 696 ±26 ng/ml; p=0.02). Melatonin: comparably to BMP-5, Melatonin stimulation resulted in aggravation of AKI in the ‘IRI+MV+Mela’ group (IRI+MV+Mela vs. IRI 872 ±34 vs. 696 ±26 ng/ml; p=0.0007). The cystatin C results are summarized in figure 1.

**Figure 1:**
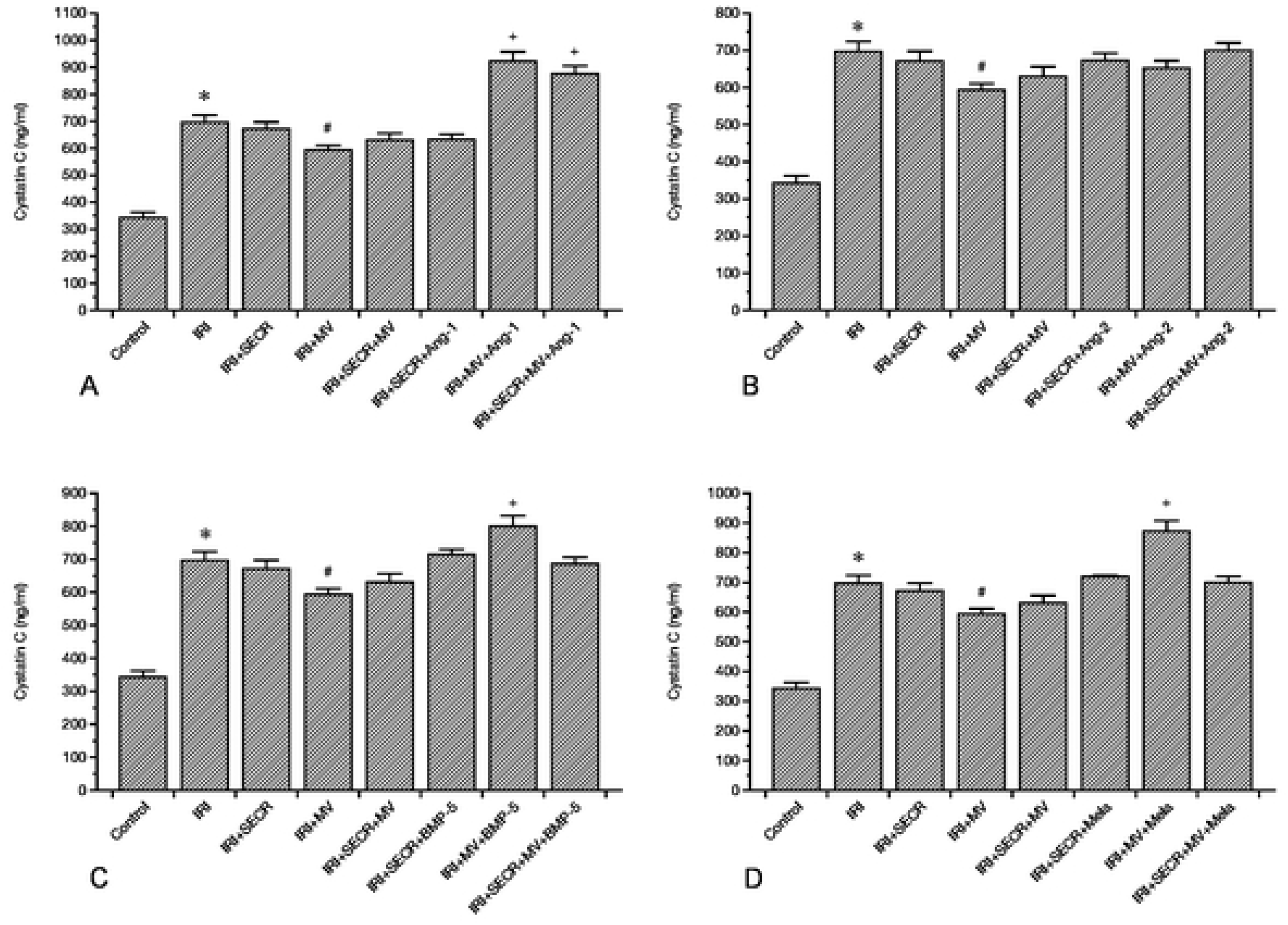
Serum Cystatin C in all control and experimental groups: The groups ‘Control’, ‘IRI’ (Ischemia Reperfusion Injury), ‘IRI+SECR’ (secretome), ‘IRI+MV’ (microvesicles), and ‘IRI+SECR+MV’ are part of every panel (A-D). A depicts the consequences of Angiopoietin-1 (Ang-1) preconditioning, B-D depict sthe effects of Angiopoietin-2 (Ang-2), melatonin (Mela), and Bone Morphogenetic Protein-5 (BMP-5) preconditioning. ✻: p-value as compared to ‘Control’ <0.05; #: p-value as compared to ‘IRI’ <0.05; +: p-value as compared to ‘IRI’ <0.05 (Results as means +/− SEM).

### Early mesenchymal transdifferentiation of endothelial cells is attenuated by native MV

AKI worsens the prognosis of respective subjects in the short- and in the long-term. The long-term prognosis critically depends on AKI-associated vascular rarefaction and interstitial fibrosis. The pathogenesis of latter is complex. However, mesenchymal transdifferentiation of endothelial cells (EndoMT) is commonly regarded as an essential fibrogenetic event [18]. We therefore aimed to evaluate endothelial expression of alpha-Smooth Muscle Actin (aSMA) early (48 hours) after ischemia. The results are given in percentages of the endothelial (CD31+) cell area (number of green pixels), covered by aSMA. In control mice, the mean percentage was 2.9 ±0.8. Post-ischemic animals (IRI) showed significantly higher values (14.9 ±2.2%; p=0.0006). Injection of the native SECR did not modulate endothelial aSMA in a significant manner (18.4 ±0.9%; p-value vs. ‘IRI’ 0.17). The administration of native MV in contrast reduced endothelial aSMA (9.3 1.2 vs. 14.9 ±2.2%; p=0.04). If native SECR and MV were injected simultaneously, 11.2 ±0.6% of the endothelial surface were covered by aSMA (p-value vs. ‘IRI’ 0.12). Regarding all other interventional groups, only one significant difference occurred as compared to the ‘IRI’ group: ‘IRI+SECR+MV+BMP-5’ showed lower endothelial aSMA abundance (8.9 ±0.9 vs. 14.9 ±2.2%; p=0.02). Figure 2 summarizes the EndoMT analyzes. In addition, we quantified interstitial matrix deposition in all groups after masson trichrome staining as published previously [14,19]. Matrix accumulation did not significantly differ between the control and any of the respective post-ischemic groups or between ‘IRI’ and one or more of the interventional groups. We avoid to mention numerical results in the text but refer to Figure 3.

**Figure 2:**
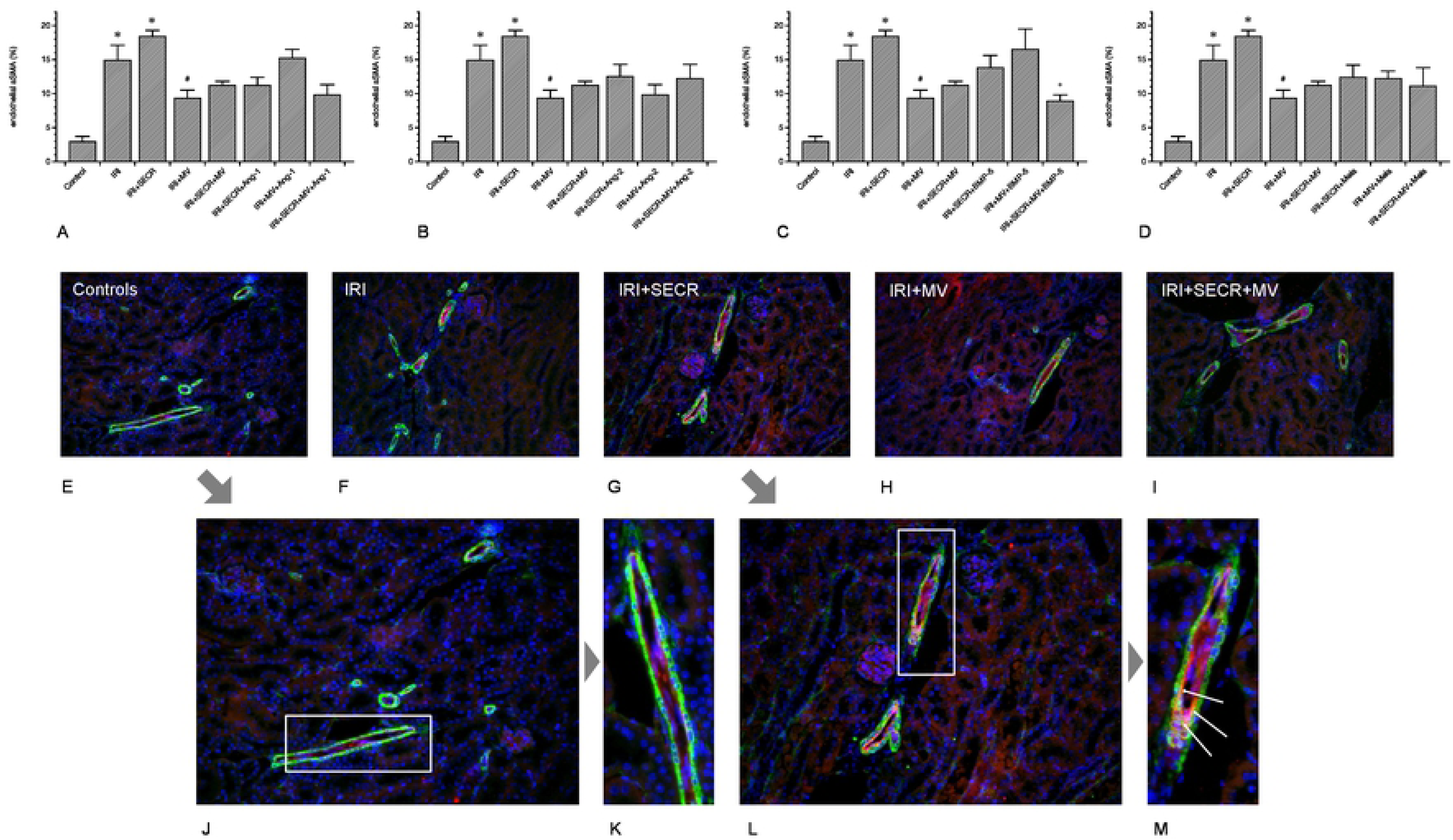
EndoMT-analyses. Comparable to figure 1, every panel (A-D) contains the results of the following groups: ‘Control’, ‘IRI’ (Ischemia Reperfusion Injury), ‘IRI+SECR’ (secretome), ‘IRI+MV’ (microvesicles), and ‘IRI+SECR+MV, respectively. E-I show representative images of small arteries / arterioles, stained for CD31 (red) and aSMA (green). The nuclei appear in blue. J-M are magnified regions of E (J+K) and G (L+M). Magnifications: E-I ×40; J and L ∼×80; K and M ∼×120. ✻: p-value as compared to ‘Control’ <0.05; # and +: p-value as compared to ‘IRI’ <0.05 (Results as means +/− SEM).

**Figure 3:**
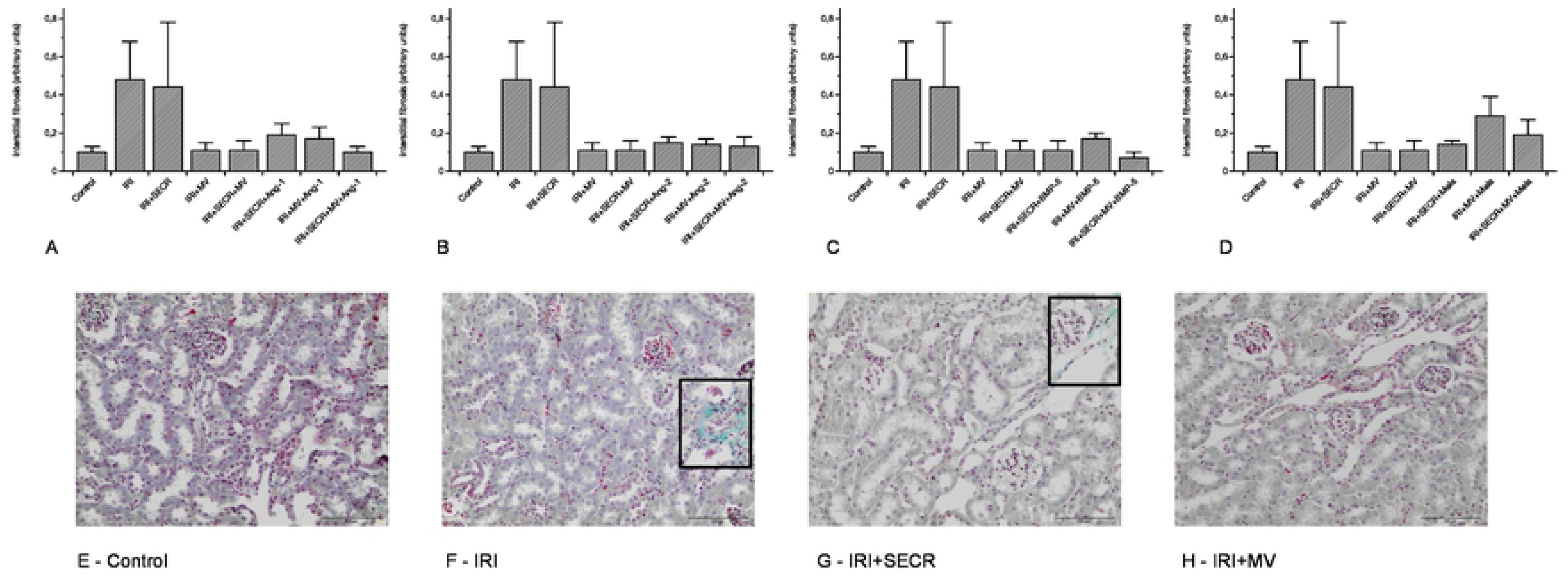
Interstitial fibrosis, evaluated after masson trichrome staining. A-D show the results of the respective preconditioning groups, always compared with ‘Control’, ‘IRI’ (Ischemia Reperfusion Injury), ‘IRI+SECR’ (secretome), ‘IRI+MV’ (microvesicles), and ‘IRI+SECR+MV, respectively. Interstitial matrix accumulation did not significantly differ at all between any of the groups. E-H (magnification ×40) show representative images of four group (labelled). The rectangels in F and G surround areas of matrix deposition.

### MV administration is associated with early peritubular capillary rarefaction

Besides interstitial fibrosis, the loss of peritubular capillaries (peritubular capillary rarefaction) is a characteristic hallmark of progressive kidney disease. It frequently results from acute renal ischemia [20]. We evaluated the density of peritubular capillaries (PTCD) by relating the CD31+ area to the total view field. We exclusively analyzed the renal cortex and did not include glomeruli. Our analyzes revealed comparable capillary densities in the control and in almost any of the interventional groups (Figure 4). The only difference occurred in post-ischemic animals that received either native MV alone or MV and SECR from unstimulated cells in combination. In these two groups the PTCD was lower as compared to even control animals, respectively (MV 0.48 ±0.1% and 0.51 ±0.1% vs. 1.2 ±0.3%; p=0.04 and p=0.03). Figure 4 shows the data in detail.

**Figure 4:**
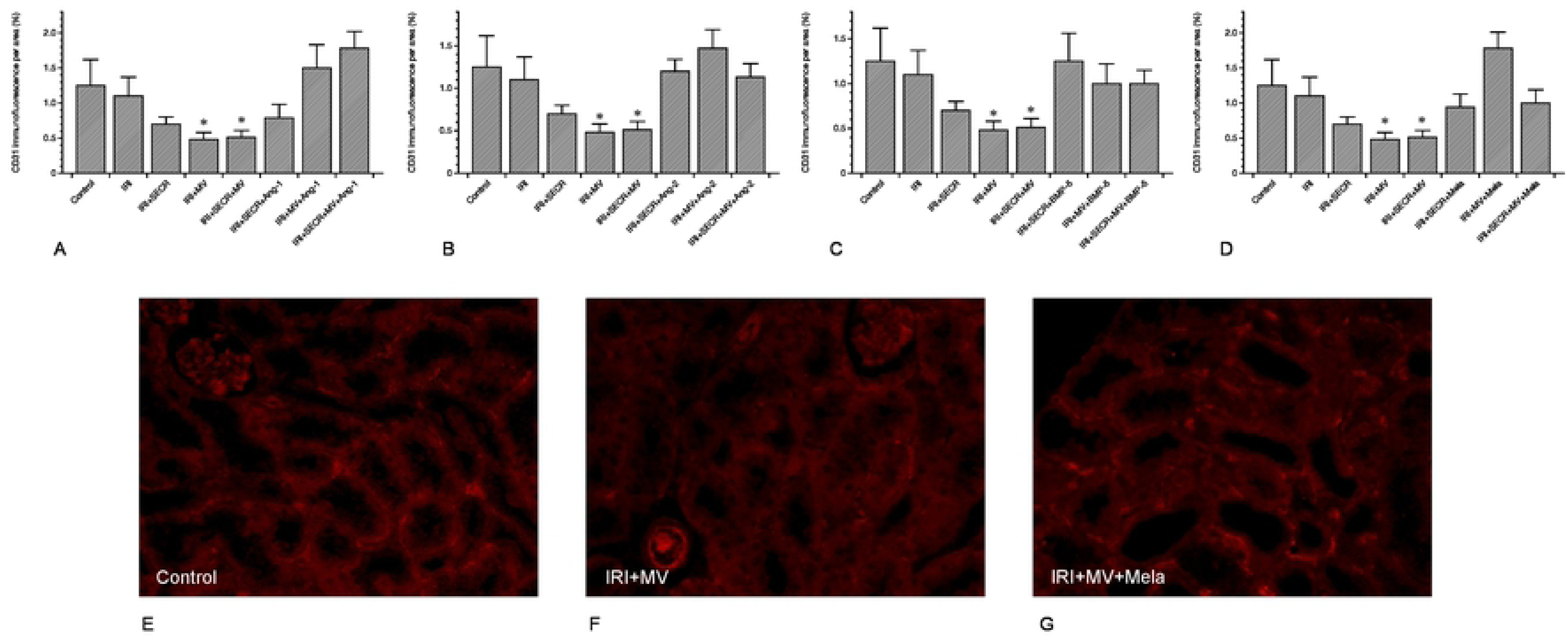
peritubular capillary density (PTCD) of all groups. The PTCD surprisingly decreased if MV from native PACs were administered (either alone or in combination with the secretome). Other significant differences were missing. A-D depicts the respective results of all preconditioning groups (A – Ang-1; B – Ang-2; C – BMP-5; D – melatonine). E-G show representative images of three groups (labelled) (magnification in E-G ×40;✻: p-value as compared to ‘IRI’ <0.05; Results as means +/− SEM).

### Endothelial alpha-Tubulin abundances do not vary early after IRI

We previously discussed reduced endothelial alpha-Tubulin (aT) expression to reflect an endogenous process of self-defense [14]. It needs however to be mentioned that respective analyzes were performed several weeks after IRI (1, 4, and 6 weeks). In the current study, endothelial aT was exclusively quantified at 48 hours after ischemia. Endothelial alpha-Tubulin did not significantly differ between the control and any of the respective post-ischemic groups or between ‘IRI’ and one or more of the interventional groups (comparably to interstitial matrix accumulation - see above). We once again avoid to mention numerical results in the text and refer to Figure 5.

**Figure 5:**
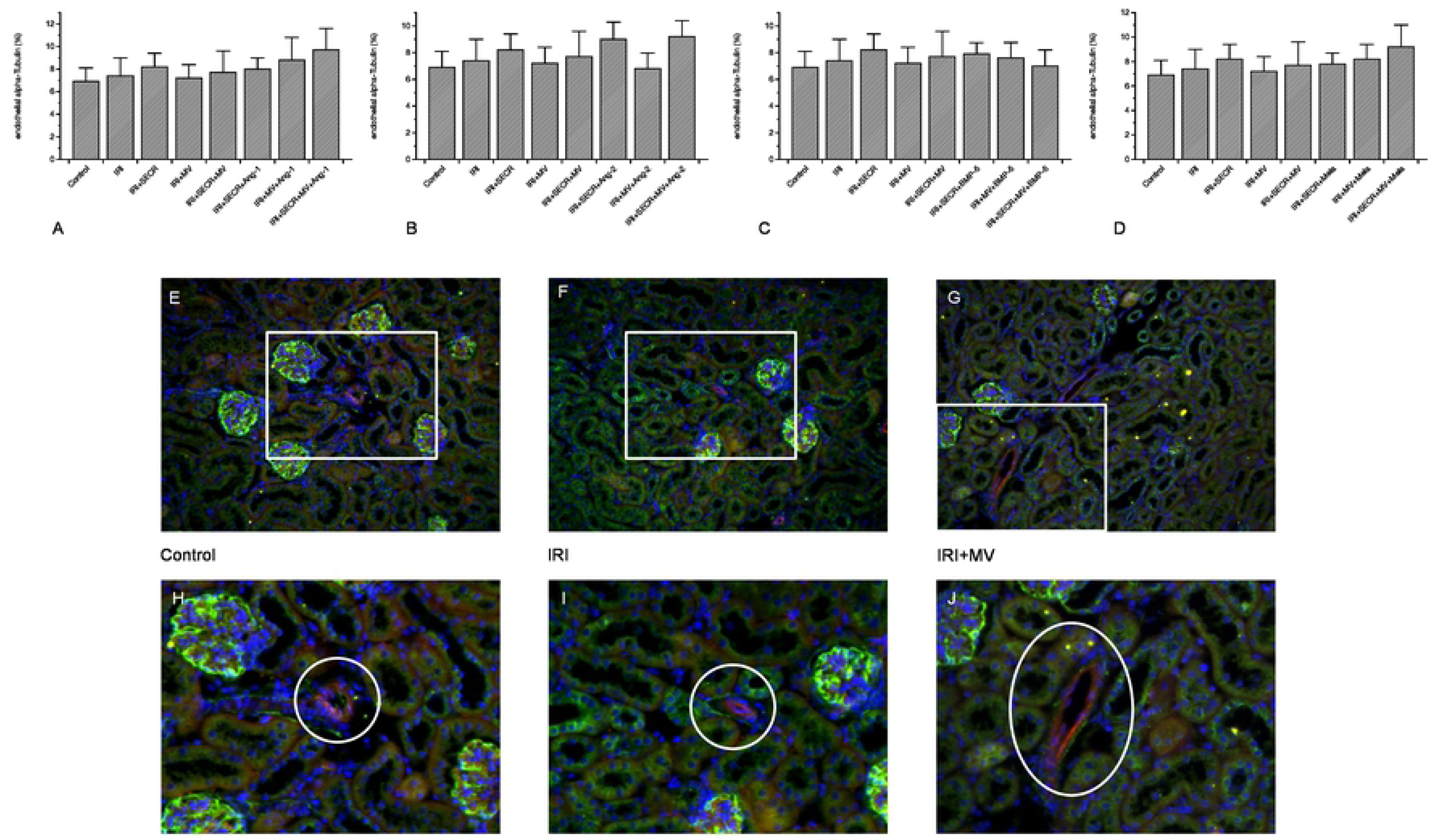
endothelial alpha-Tubulin expression in all groups. We did not detect any significant difference at all. A-D: illustration of the numerical results (A – Ang-1; B – Ang-2; C – BMP-5; D – melatonine). E-F: representative images of three groups (labelled, magnification ×40); the white rectangles surround the magnified areas of H-J (magnification ×160); the white circles in H-J indicate the vessel of interest (Results as means +/− SEM).

### Proteomic analysis of the PAC-derived secretome

To better explore the protective role of PACs we investigated the alteration of the PAC-secretome under native and stimulated conditions. For this purpose, a deep proteomic investigation was performed. We generated secretomes from control, BMP-5 and Ang-2 treated PACs. The procedure including 1D-SDS-PAGE separation followed by in gel digestion and LC-MS/MS with spectral account quantification was carried out for each sample. In order to obtain a quick overview on the composition of the secretome, we plotted fold-changes versus significance to generate volcano plots. This allow a quick identification of secretome proteins that were significantly (p<0.05) altered in their levels upon stimulation. Comparative presentation of the mass spectrometry data in Supplemental Table 1 allowed an overview on the large secretome alteration and the identification of a number of proteins that were only identified in the secretomes of Ang-2 or BMP-5 treated PACs.

As far as Ang-2 stimulated proteins were considered alone, the resulting interactome was highly complex (Figure 6A). The (expanded) pathway analysis however revealed a group of proteins that could be divided into five main categories: extracellular matrix organization, innate immune system, neutrophil degranulation, metabolism and immune system. Network analysis of the proteins, which secretion was inhibited upon Ang-2 treatment resulted in a less complex secretome with proteins involved in the coagulation cascade and platelet activation (Figure 6B). BMP-5 treatment of PACs resulted in diminution of a large number of proteins that are principally involved in almost the same pathways (immune system, innate immune system, hemostasis, platelet degranulation, platelet activation, signaling and aggregation) as found in the Ang-2 analyzes (Figure 6C). The comparison of both, the Ang-2 and BMP-5 secretomes showed up-regulation of proteins involved in joint pathways (Figure 6D).

**Figure 6:**
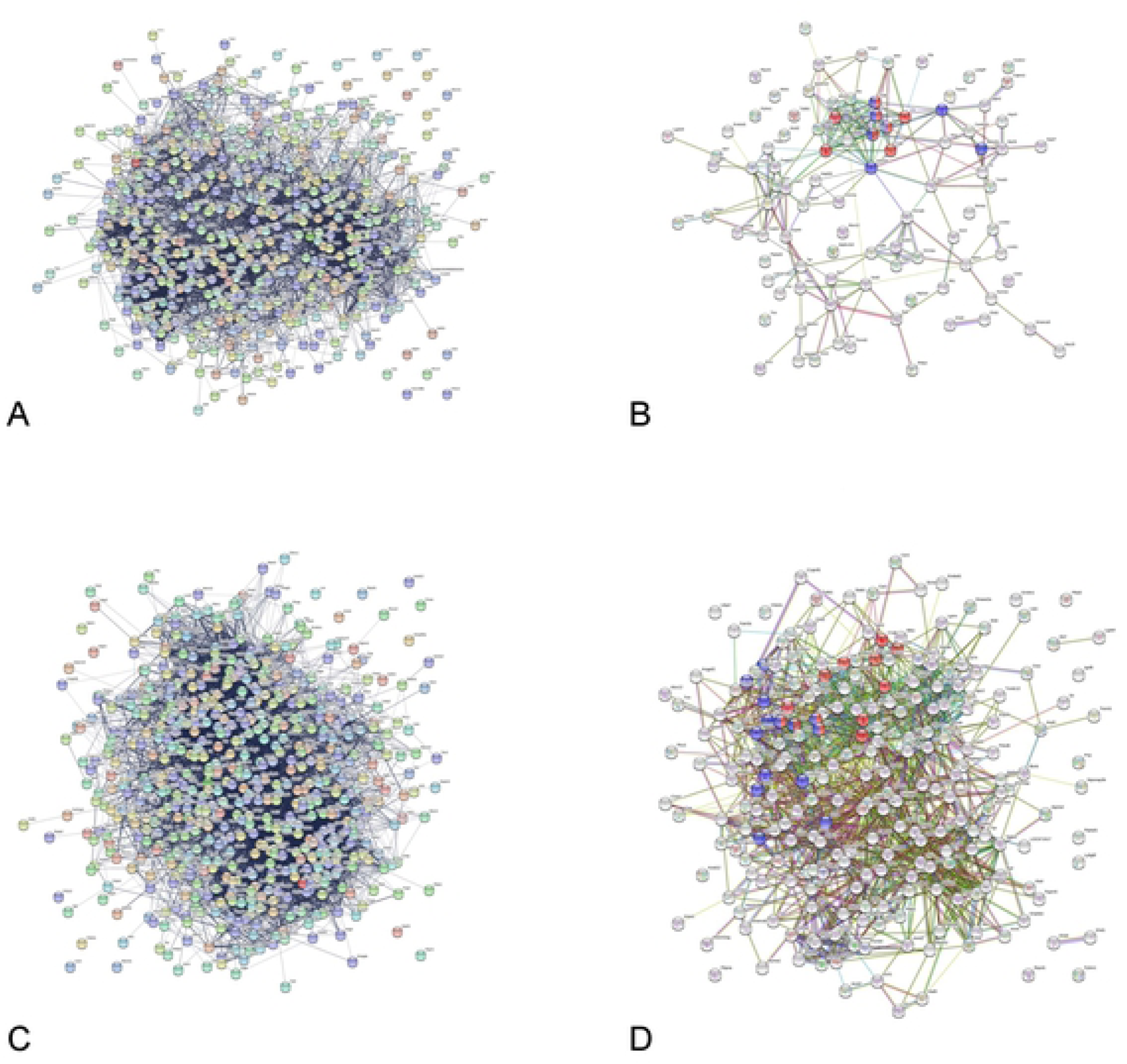
Protein interaction networks between the proteins found to be differentially secreted upon treatments were generated using the protein networks software String (https://string-db.org). A: Interaction network of the secretome proteins that were more than two-fold secreted in Ang-2 treated cells compared to controls (ratio Agn-2/Ctr > 2); B: Interaction network of proteins found to be down-regulated (ratio Agn-2/Ctr < 0.5) under Ang-2 treatment compared to control. Interestingly the interaction network showed that Ang-2 treatment resulted in decreased secretion of proteins involved in the coagulation cascade and in platelet activation (red and blue colored). C: Interaction network of proteins, which level was down-regulated (ratio BMP5/Ctr < 0.5) in secretome of BMP5 treated cells; D: Interaction network of proteins highly secreted upon Ang-2 treatment compared to BMP5 treatment (ratio Ang-2/BMP5 > 2). The network revealed that Ang-2 treatment resulted in significant alteration of EPC-secretome.

## Discussion

The most intruiging findings in the current study were the exclusive improvement of post-ischemic kidney function after injecting MV from native cells and the complete lack of stimulatory effects of any of the four mediators on either MV or SECR. Two conclusions need to be drawn: MV substantially participate in mediating AKI protection, but cell-derived extracellular components are most likely not solely responsible for improving kidney dysfunction in acute situations.

In the past, we showed AKI protective effects of injected PACs in numerous experimental studies [8–10]. All protocols employed so far utilized intact cells, either without or after pharmacological preconditioning. Over the years we identified five distinct mediators that substantially improved the cells’ therapeutic competence in AKI [8,15–17,21]. The current study, which avoided the injection of living cells did not confirm any of these observations. Thus, we must conclude that the mere presence of cells within the kidney is important for PAC-associated kidney protection, particularly under circumstances when preconditioning protocols are being applied. The field of PAC(EPC)-mediated AKI protection substantially benefited from studies by Cantaluppi and colleagues [12] who identified the critical role of cell-derived microvesicles (MV) in IRI. The authors did not only prove that AKI may be prevented by injecting MV alone but they also extracted certain microRNA molecules from the vesicular fraction. By RNA depletion prior to MV administration, protective effects of the vesicles vanished. The discussion about cellular mechanisms responsible for tissue protection not only in AKI and not only regarding PACs has been going on for a long time. In 2003, Rehman et al. showed that supernatant from cultured EPCs are significantly enriched by angiogenic growth factors such as vascular endothelial growth factor, hepatocyte growth factor, granulocyte colony-stimulating factor, and by granulocyte-macrophage colony-stimulating factor [13]. Since then, cell effects were thought to substantially occur in an indirect manner. In subsequent years, proteomic analyzes revealed additional humoral mediators including thymidine phosphorylase, matrix metalloproteinase 9, IL-8, pre-B cell enhancing factor, and macrophage migration inhibitory factor [22,23]. Thus, the concept of the ‘secretome’ evolved. The profound modulatory role of the secretome was highlighted by studies of Goligorsky’s group. In a 2015 published article, EPC-/MSC(Mesenchymal Stem Cell)-derived cytokines substantially influenced the phenotype and biological behavior of macrophages [24]. It was concluded that the secretome partly mediates protection from LPS-induced AKI. Later studies showed that endothelium-derived humoral factors are also involved in renal fibrogenesis [25]. In the 2015 published article [24] one essential problem related to any cell-based therapeutic approach in AKI was emphasized: the problem of cell delivery. Therefore, MV appear as promising option at first sight, although the problem of ex vivo generation of such highly complex structures would have to be solved. In addition, our analyses did not exclusively show beneficial end point effects of MV. For instance, cystatin C levels increased and anti-mesenchymal effects vanished if MV from preconditioned cells (Ang-1, Mela, BMP-5) were applied. Finally, MV from native PACs significantly affected the vascular structure despite the fact that excretory function was improved.

In the last paragraph will would like to discuss our proteomic data. Only few studies evaluated the role and composition of the secretome derived from so-called Endothelial Progenitor Cells (here: PACs) in the past. In one study, published in 2015 [24], hydrogel-embedded EPCs were subcutaneously implanted in LPS-treated mice, resulting in AKI protection under specific circumstances. While the study addressed systemic cytokine levels in the animals, the authors did not acquire data on the secretome per se. It was however concluded that paracrinic actions of (embedded) EPCs most likely mediate biological cell effects. The possibility of MV-dependent activity was not discussed at all. A 2018 published investigation analyzed the EPC secretome in the context of oligodendrocyte repair [26]. The authors performed a detailed proteomic study and identified a heterogenous group of proteins excreted by the cells. Among those were angiogenin, stromal derived factor-1 (SDF-1), platelet-derived growth factor, vascular endothelial growth factor-B (VEGF-B), and several matrix metalloproteinases (MMP). Thus, they detected mediators with substantial proangiogenic activity. The third study that needs to be mentioned focused on EPC secretome-enriched nanoparticles in vascular repair [27]. It was documented that hypoxia stimulated the production of secretome proteins with angiogenic properties. Contrasting to these reports, our analyses revealed enrichement of mediators involved in the coagulation cascade and platelet activation. Significant stimulatory effects on angiogenic proteins did not become apparent. The discrepant findings possibly result from the artificial conditions that were used in our investigation. Felice and colleagues [27] exposed the cells to hypoxic conditions while we subjected PACs to specific mediators that were shown to significantly increase the AKI protective activity of intact cells [8,15–17]. Regarding the fact that our study failed to show any beneficial effects of the secretome at all, the data substantially indicate that renoprotective effects of whole cells most likely require the mere presence of intact PACs in the kidney, especially if preconditioning protocols are in use.

Together, our data show that PAC-mediated AKI protection substantially depends on the availability of cell-derived EV. However, since previous data showed improved AKI-protection by PACs after cell preconditioning with certain mediators (Ang-1 and −2, melatonine, BMP-5), other than exclusively vesicle-dependent mechanisms must be involved in PAC-mediated AKI protection.

## Material and methods

### Animals models

As in previous animal-based studies, all protocols were performed according to the guidelines of the German Institute of Health Guide for the Care and Use of Laboratory Animals and approved by the Institutional Animal Care and Use Committee of the University of Göttingen. For all experiments, we employed male C57/Bl6N mice (8-12 weeks old). Mice were bred in the local animal facility of the Göttingen University Hospital. Animals were separately caged with a 12:12-h light-dark cycle and had free access to water and chow throughout the study.

### Surgical procedures

Anaesthesia was performed with a solution of 300 μl 6 mg/100 g ketamine hydrochloride plus 0.77 mg/100 of xylazine hydrochloride, applied intraperitoneally. Mice were placed on a heated surgical pad during the whole procedure. Rectal temperature was permanently maintained at 37°C. The abdominal cavity was opened by a 1.5-cm midlaparotomy. Both kidneys were exposed and clamping of the renal pedicles was performed with microserrefines (Fine Science Tools, Forster City, USA) for 40 minutes, respectively. The clamps were released and a constant volume of MV or SECR containing buffer / medium (Isolation of cell-derived microvesicles (MV) and secretome (SECR) from native and preconditioned PACs) was injected into the tail vein and thus, into the systemic circulation. The abdominal incision was closed with a 4-0 suture and surgical staples. In each experimental group 10 animals were analyzed. Animals were sacrificed 48 hours later. Euthanization was performed by injecting the threefold dose of anaethesia, followed by dissecting the diaphragm.

### Isolation of cell-derived microvesicles (MV) and secretome (SECR) from native and preconditioned PACs

Both components, PAC-derived microvesicles (MV) and the secretome (SECR) were isolated from syngeneic murine PACs. The cells were isolated and expanded as described previously in detail [14]. However, we will briefly describe the procedure again. Mouse mononuclear cells (MNCs) were enriched by density gradient centrifugation using Biocoll solution (Biochrom, Berlin, Germany) from peripheral blood and spleen cell extracts. The reason for collecting cells from the two compartments was the intention to maximize the total number of cells available for injection. Mononuclear cells were mixed and, differing to previous protocols in which 4×106 cells were used [14], 25×106 cells were plated on 6-well culture dishes coated with human fibronectin (Sigma, St Louis, MO) and maintained in endothelial cell medium-2 (EGM-2 -Clonetics, Lonza, Walkersville, MD, USA) supplemented with endothelial growth medium (EGM) Single-Quots containing 5% FCS. Four to 5 days later, cultured PACs were identified by the uptake of DiI-labeled acetylated low density lipoprotein (acLDL) (Invitrogen, Carlsbad, CA, USA) and simultaneous binding of FITC-labeled BS-1 lectin (BS-1) (Sigma Diagnostics, St. Louis, MO). SECR: at the day of surgery (5-6 days after initial cell seeding), PACs were incubated with one of the following substances: Angiopoietin-1 (250 ng/mL); Angiopoietin-2 (250 ng/mL); Bone Morphogenetic Protein-5 (BMP-5 – 100 ng/mL); melatonin (5 µMol/L).

Incubations were performed for 1 hour at 37°C, the mediators were applied in fresh EGM-2, respectively. Supernatants were collected and filtered (pore size 0.8 µm). Filtrates were subsequently centrifuged for 6 minutes at 13,000 rpm auf (Vivaspin 500 100,000 MWCO - Satorius VS0142). For SECR injection experiments, every mouse received 100 µL per tail vein injection. MV: after supernatant filtration (pore size 0.8 µm), cell-derived microvesicles were collected using the ExoEasy Kit (Qiagen 76064) according to the manufacturer’s protocol. Finally, a total volume of MV-containing eluate was applied in an individual mouse per tail vein injection.

### Histology and immunoflurescence staining of tissue sections

The following methodical approaches were used for the quantification of fibrosis (masson trichrome) and of endothelial alpha-Smooth Muscle Actin (aSMA) or alpha-Tubulin (aT) expression. All analyses were performed using ImageJ software. Fibrosis: the cortical area of every kidney was documented, areas not being covered by tissue were not considered. The total area of analysis was quantified by the total number of pixels. All pixels within a particular colour range (here: green) were quantified and related to the total pixel number of interest. Results were given as percent of the total number of image pixels. Co-localization analysis (CD31 in combination with aSMA or aT): in every kidney section, at least three different small blood vessels were evaluated in an individual manner. The endothelial surface was quantified by counting the total number of red pixels (CD31). Yellow areas were also quantified, according to a defined color range. The colour yellow reflected the respective marker of interest in endothelial cells, the absolute number of yellow pixels was related to the absolute number of red pixels, respectively (results in percent). Kidney fibrosis was examined in formalin fixated, paraffin-embedded tissue sections after masson trichrome staining. Endothelial expression of aSMA and aT were also analyzed in formalin fixated, paraffin-embedded tissue sections after deparaffinization, followed by incubation in 3% H_2_O_2_ for 10 minutes. After citrate-buffer treatment (microwave, 5 times 3 minutes, pH 6.0) sections were stained with rat anti-mouse CD31 (PECAM-1 - CloneSZ31, Dianova), and rabbit anti-Smooth Muscle Actin (EMELCA) or mouse anti-alpha Tubulin (abcam - ab24610) for primary incubation and with Alexa Fluor 488 goat anti-rabbit IgG (Dianova) and Alexa Fluor 594 goat anti-rat IgG (Dianova) or Alexa Fluor 488 anti-mouse IgG (Dianova) for secondary incubation, respectively. Primary incubation was performed overnight at 4°C while secondary incubation was performed for 1 hour at room temperature. To visualize the nuclei, tissue sections were counterstained with DAPI.

### Quantification of peritubular capillary density (PTCD)

For PTCD quantification, exclusively cortical areas not containing glomeruli were documented. The total area of interest was evaluated for the absolute number of pixels. After CD31 staining (*Histology and immunoflurescence staining of tissue* sections) the colour red was quantified as described above and related to the total area of interest (red pixels as percentage of all pixels per area).

### Quantification of serum Cystatin C

Serum Cystatin C levels were quantified using a commercially available kit (BioVendor, RD291009200R) according to the manufacturer’s instructions.

### PAC isolation and treatment for proteomic analyses

PACs from wildtype mice blood and spleen were isolated, expanded over 5-6 days of culture in EGM-2-medium (Lonza, CC-3156, CC-4176). Medium was aspirated and replaced by pre-warmed 10 mL serum-free EGM-2-medium per 10 cm/dish. The serum-free EGM-2-medium was replaced by fresh medium 3 times, each incubation period lasting for 120 minutes in the presence of CO_2_. Finally, medium was replaced by FCS free EGM-2 containing either Ang-2 (200 ng/ml) or BMP-5 (100 ng/ml). The treatment was carried out for 48 hours in an CO_2_ incubator at 37°C. Subsequently conditioned medium was collected, centrifuged at 300g for 5 minutes at 4°C, supernatants collected and stored at −80°C.

### Protein precipitation and concentration estimation

To enrich the secretome proteins, the supernatants from control and treated PACs (10 ml each) were concentrated separately to 2 ml with a Vivaspin 20 Ultrafiltration Unit (Sartorius Göttingen, Germany). The resulting samples were then subjected to protein precipitation to reduce the volume and enrich the proteins. The precipitation was carried out by adding 3 volumes of ice-cold acetone containing 10% methanol and incubating overnight at −20°C. Precipitated proteins were pelleted by centrifugation at 12,000g for 45 min at 4°C. The pellets were dried and resolved in urea buffer (30 mM Tris-HCl pH 8.5, 9.5 M urea, 2% CHAPS). The protein concentration was determined according to the Bradford method using BSA as the calibrator.

### SDS-PAGE, tryptic in gel digestion and mass spectrometric analysis of the PAC-derived secretome

The protein extracts from control sample and the different treatments were separated in 1D-SDS-PAGE. The gels were stained with Coomassie blue and the visualized protein lanes were excised in 20 band section each and subjected to in gel digestion with trypsin (12.5 ng/µl). The resulting tryptic digests were extracted and analyzed using the Thermo Scientific Q Exactive. All MS/MS samples were analyzed using Mascot (Matrix Science, London, UK; version 2.4.1). Mascot was searched with a fragment ion mass tolerance of 0.020 Da and a parent ion tolerance of 10.0 PPM. Scaffold (version Scaffold_4.8.9), Proteome Software Inc., Portland, OR) was used to validate MS/MS based peptide and protein identifications. The software normalizes the MS/MS data between samples. This allows us to compare abundances of a protein between samples. The normalization scheme used works for the common experimental situation where individual proteins may be up-regulated or down-regulated, but the total amount of all proteins in each sample is about the same like it was the case in our experiments. Normalization is done on the MS sample level. The normalization method that Scaffold uses is to sum the “Unweighted Spectrum Counts” for each MS sample. These sums are then scaled so that they are all the same. The scaling factor for each sample is then applied to each protein group and adjusts its “Unweighted Spectrum Count” to a normalized “Quantitative Value”.

### Bioinformatic Analyses

To examine potential protein function categories and pathways of significantly regulated proteins, we performed bioinformatics analysis using the following public protein software: DAVID Functional Annotation Bioinformatics Microarray Analysis (http://david.abcc.ncifcrf.gov/), the protein-protein interaction network software String 10.5 (https://string-db.org), and the pathway analysis software Kegg (https://www.genome.jp/kegg/).

### Statistical analyses

All results are given an mean +/− SEM. The means of two groups were compared using the student’s ttest. Differences between three or more groups were analyzed with the ANOVA test. Significance was postulated if the p-value was below 0.05.

## Author contributions

### Conceptualization

Hassan Dihazi, Daniel Patschan

### Data curation

Katrin Schwarze

### Formal analysis

Hassan Dihazi, Katrin Schwarze, Daniel Patschan

### Funding acquisition

Daniel Patschan

### Investigation

Hassan Dihazi, Katrin Schwarze, Susann Patschan, Daniel Patschan

### Methodology

Hassan Dihazi, Daniel Patschan

### Project administration

Katrin Schwarze

### Resources

Hassan Dihazi, Katrin Schwarze, Susann Patschan, Daniel Patschan

### Software

Hassan Dihazi, Daniel Patschan

### Supervision

Hassan Dihazi, Daniel Patschan

### Validation

Hassan Dihazi, Susann Patschan, Daniel Patschan

### Visualization

Hassan Dihazi, Daniel Patschan

### Writing – original draft

Daniel Patschan

### Writing – review & editing

all authors

